# Aging with TBI vs. Aging: 6-month temporal profiles for neuropathology and astrocyte activation converge in behaviorally relevant thalamocortical circuitry of male and female rats

**DOI:** 10.1101/2023.02.06.527058

**Authors:** Zackary Sabetta, Gokul Krishna, Tala Curry, P. David Adelson, Theresa Currier Thomas

## Abstract

Traumatic brain injury (TBI) manifests late-onset and persisting clinical symptoms with implications for sex differences and increased risk for the development of age-related neurodegenerative diseases. Few studies have evaluated chronic temporal profiles of neuronal and glial pathology that include sex as a biological variable. After experimental diffuse TBI, late-onset and persisting somatosensory hypersensitivity to whisker stimulation develops at one-month post-injury and persists to at least two months post-injury in male rats, providing an *in vivo* model to evaluate the temporal profile of pathology responsible for morbidity. Whisker somatosensation is dependent on signaling through the thalamocortical relays of the whisker barrel circuit made up of glutamatergic projections between the ventral posteromedial nucleus of the thalamus (VPM) and primary somatosensory barrel cortex (S1BF) with inhibitory (GABA) innervation from the thalamic reticular nucleus (TRN) to the VPM. To evaluate the temporal profiles of pathology, male and female Sprague Dawley rats (*n* = 5-6/group) were subjected to sham surgery or midline fluid percussion injury (FPI). At 7-, 56-, and 168-days post-injury (DPI), brains were processed for amino-cupric silver stain and glial fibrillary acidic protein (GFAP) immunoreactivity, where pixel density of staining was quantified to determine the temporal profile of neuropathology and astrocyte activation in the VPM, S1BF, and TRN. FPI induced significant neuropathology in all brain regions at 7 DPI. At 168 DPI, neuropathology remained significantly elevated in the VPM and TRN, but returned to sham levels in the S1BF. GFAP immunoreactivity was increased as a function of FPI and DPI, with an FPI × DPI interaction in all regions and an FPI × Sex interaction in the S1BF. The interactions were driven by increased GFAP immunoreactivity in shams over time in the VPM and TRN. In the S1BF, GFAP immunoreactivity increased at 7 DPI and declined to age-matched sham levels by 168 DPI, while GFAP immunoreactivity in shams significantly increased between 7 and 168 days. The FPI × Sex interaction was driven by an overall greater level of GFAP immunoreactivity in FPI males compared to FPI females. Increased GFAP immunoreactivity was associated with an increased number of GFAP-positive soma, predominantly at 7 DPI. Overall, these findings indicate that FPI, time post-injury, sex, region, and aging with injury differentially contribute to chronic changes in neuronal pathology and astrocyte activation after diffuse brain injury. Thus, our results highlight distinct patterns of pathological alterations associated with the development and persistence of morbidity that supports chronic neuropathology, especially within the thalamus. Further, data indicate a convergence between TBI-induced and age-related pathology where further investigation may reveal a role for divergent astrocytic phenotypes associated with increased risk for neurodegenerative diseases.

## INTRODUCTION

Traumatic brain injury (TBI) continues to be one of the most complex brain injuries with long-term consequences that can lead to persistent and late-onset symptoms and increased risk for developing age-associated neurodegenerative disorders [1]. Mild TBI or concussive head injury (Glasgow Coma Scale score ≥13) constitutes over 80% of all TBI cases globally and often leads to heterogenous clinical symptoms that can be increasingly difficult for patients to manage [2–4]. Several recent studies indicate that mild TBI patient populations tend to have a greater impact on the overall economic burden due to the sheer number of patients with persisting symptoms that impede return to work and require subsequent healthcare [2, 5]. Furthermore, while aging is an associated risk factor for neurodegenerative conditions, little is known regarding how the brain compensates for aging with TBI, where TBI is implicated in accelerating brain aging or exacerbating neurodegenerative pathology [6–8]. Moreover, while reported TBI is higher in males than females [9], clinical evidence is conflicting concerning sex-dependent recovery after TBI where previous studies indicate sex-dependent outcomes and that females often display more severe co-morbid symptoms [10–12]. On the other hand, males more commonly report chronic symptoms such as aggression, sleep disturbances, and substance abuse [13]. With the recently increased recognition of TBI in domestic violence [11, 14], which primarily targets women and children, a better understanding of the sex-specific impact of TBI on pathological outcomes is required.

Post-TBI secondary cascades show distinct and dynamic pathological features at acute, subacute, and chronic stages [15]. Brain-injury mediated events involve maladaptive circuit reorganization with development of a broad spectrum of chronic morbidities and it is becoming increasingly evident that glial cells are important players in the pathological process. Astrocytes respond rapidly as sentinels undergoing expressive changes following TBI to influence neuroinflammation, neurotransmission, neuroplasticity, neuroprotection, neurotoxicity, and play a central role in blood-brain barrier (BBB) integrity [16, 17]. Reactive astrocytes are generally characterized by the upregulation of glial fibrillary acid protein (GFAP) associated with morphological alterations (i.e., hypertrophy) and astrocyte proliferation. Reactive astrocytes in response to TBI pathology are associated with chronic neurodegeneration and neuronal dysfunction [18], amplified release of pro-inflammatory mediators [17], and altered ion influx [17, 19, 20] which may contribute to the post-TBI persistence and progression of neurodegeneration [21]. Further, an inflammatory astrocytic phenotype has been identified as a function of age [22] and is associated with loss of proteostasis, mitochondrial dysfunction, and altered cellular communication [23, 24]. However, there is little information on the temporal dynamics of TBI-induced astrocyte activation or how aging with TBI and normal aging can sex-specifically influence the magnitude and duration of astrocytic changes.

Among the common clinical morbidities resulting from TBI, sensory hypersensitivity is reported in auditory, visual, and tactile sensations for several years after the initial insult [25, 26]. Following experimental diffuse TBI, circuit disruption and consequent reorganization lead to changes in circuit function that correspond with the development of late-onset sensory hypersensitivity to whisker stimulation in both sexes, and this effect, was found to be lower in females than males [14, 27, 28]. Whiskers are the primary sensory modality for rodents and behaviorally relevant somatosensory information is relayed through neuronal circuits distributed across multiple brain regions, where the development and persistence of post-TBI symptoms can be related to a single relay or the culmination of dysfunction in several relays. Tactile sensory input from each whisker is communicated to discreet corresponding circuit relays in the midbrain, primary ventral posteromedial nucleus of the thalamus (VPM), and somatosensory barrel cortex (S1BF) through the lemniscal pathway. The rodent whisker barrel circuit (WBC) is predominantly glutamatergic, with inhibitory innervation from the thalamic reticular nucleus (TRN) that largely controls the input and output of the circuit [29]. Hypersensitivity is indicated by an atypical behavioral response to whisker stimulation associated with changes in movement, evasion, body position, breathing, whisker position, whisking, and grooming [27]. These behavioral changes are concomitant with sex-dependent changes in neurotransmission, increased plasma corticosterone to whisker stimulation, and chronic disruption of the hypothalamic-pituitary-adrenal axis [14, 27, 30–32]. These late-onset and persisting sensory symptoms indicate sensory processing deficits where the temporal assessment of underlying pathophysiology can reveal translationally relevant mechanisms important for enhancing current clinical protocols. In the present study, we quantify biomarkers of neurodegeneration and astrocyte reactivity out to 6 months post-injury in the relays of the WBC to fill the knowledge gap of injury-, sex-, region-, and time-dependent pathology associated with the manifestation and persistence of post-TBI symptoms. Our study provides evidence of chronic neuropathological and astrocytic changes after a single diffuse TBI in both sexes as well as age and injury-related interactions.

## MATERIALS AND METHODS

### Animals

A total of 64 young adult age-matched male and naturally cycling female Sprague-Dawley rats (~3 months old; males 367 ± 3 g and females 235 ± 1.5 g; n = 5-6/group; Envigo, Indianapolis, IN, USA) were used in these experiments and maintained in a 12:12 h light:dark cycle in a temperature and humidity-controlled room. Food (Teklad 2918) and water (Innovive, San Diego, CA, USA) were provided *ad libitum* and rats allowed to acclimate for at least one week before experiments. Group sizes were determined from a previous publication [33], where n = 5 could predict >90% power for a significant fluid percussion injury (FPI) effect in the VPM of males. A post-hoc power calculation for MANOVA was calculated for an FPI effect with GFAP and silver stains in the VPM, where n = 5/group provided >99% power with Cohen’s f values being 1.85 and 0.92, respectively (calculations using G*Power 3.1.9.7 post hoc: compute achieved power). All procedures were conducted in compliance with ARRIVE guidelines and consistent with the National Institutes of Health (NIH) Guidelines for the Care and Use of Laboratory Animals approved by the University of Arizona Institutional Animal Care and Use Committee (protocol #18-384) at the University of Arizona College of Medicine-Phoenix.

### Surgical Procedure

Midline fluid percussion injury (FPI) was carried out using an FPI device (Custom design and Fabrication, Virginia Commonwealth University, Richmond, VA) similarly to previously published methods from this laboratory [28, 34, 35]. Each cage of rats (2/cage) was randomized into either FPI or sham groups. Briefly, rats were anesthetized with isoflurane (5% in 100% oxygen with a flow rate of 0.8 L/min), weighed, heads shaved, and placed into a stereotaxic frame (Kopf Instruments, Tujunga, CA) with a nose cone that maintained 2.5% isoflurane for the duration of the procedure. The surgical site was sterilized with alternating applications of iodine and ethanol. A 4.8 mm circular craniectomy was centered on the sagittal suture and fitted with an injury hub fabricated from the female end of a 20-gauge Luer Loc needle. A stainless-steel anchoring screw was placed in the right frontal bone. The injury hub was affixed over the craniectomy using cyanoacrylate gel and methyl-methacrylate (Hygenic Corp., Akron, OH) and filled with 0.9% sterile saline. The incision was then partially sutured closed, and topical lidocaine and antibiotic ointment were applied. Rats were returned to a warmed holding cage and monitored until ambulatory.

### Injury Induction

Approximately two hours following surgical procedures, rats were re-anesthetized using 5% isoflurane in 100% oxygen for 3 minutes. The Luer Loc hub was filled with 0.9% sterile saline and attached to the male end of a fluid percussion device. After the return of a pedal withdrawal response, an injury averaging 1.8-2.0 atmospheric pressure (atm) for males and 1.7-1.9 atm for females was administered on the dura by releasing the pendulum (from 16 degrees for males and 15.5 degrees for females) onto the fluid-filled cylinder. These atmospheric pressures were predetermined in our lab and others [36]. Shams were attached to the fluid percussion device, but the pendulum was not released after a positive pedal withdrawal response. Immediately after administration of the injury, the forearm fencing response was recorded for injured animals, and the injury hub was removed *en bloc*. Brain-injured rats were monitored for the presence of apnea and the return of righting reflex [37, 38]. The righting reflex time is the total time from initial impact until the rat spontaneously rights itself from a supine position to a prone position. Inclusion criteria required that injured rats have a righting reflex time ranging from 6-10 minutes and a fencing response. Rats were re-anesthetized for 2 minutes to inspect the injury site for hematoma, herniation, and dural integrity. The injury site was then stapled closed (BD AutoClip^™^,9mm), and topical lidocaine and antibiotic ointment were applied to minimize pain and discomfort. Rats were then placed in a clean, warmed holding cage and monitored for at least one hour following injury or sham surgery before being placed in a new, clean cage with bedding and returned to the vivarium housing room, where post-operative evaluations continued for 4 days post-injury/surgery. Post-operative monitoring included the appearance of incision, monitoring of behavior, body weight measurements, and a pain scale evaluation. Rats were euthanized (and therefore excluded) if they lost more than 15% of their body weight or presented with chronic pain symptoms, as described by the American Association for Accreditation of Laboratory Animal Care (AAALAC). No animals were excluded based on these conditions. Wound clips were removed Rats were pair-housed according to injury status and sex, remaining with the same cage mates throughout the duration of the study.

### Brain Collection

Rats did not undergo any behavior tasks; environments were minimized for potential stressors, and acute stress responses were controlled by carrying out procedures within fixed durations to prevent confounding variables that could potentially influence neuroplasticity and other neurophysiological properties. Brains were collected at 7, 56, and 168 DPI, between 07:00 and 11:00 AM to minimize any impact of diurnal variations. On the day of sacrifice, an assigned cage retriever carried a timer which was started as soon as they entered the home cage room. One cage of rats was retrieved at a time and brought directly to the necropsy suite, where both rats were immediately placed in the induction chamber that had been pre-filled with isoflurane (~30-45 s) for two minutes (5% isoflurane at 2.5 oxygen flow rate) [39]. Two teams were available (one for each rat), where rats were immediately removed from the induction chamber, weighed, rapidly decapitated, and brains were removed. The total time between cage room disruption and brain extraction was 190-220 seconds. All efforts were made to minimize interactions with the retriever and necropsy teams to prevent cross-contamination of odors between the necropsy suite and cage room. Extracted brains were rinsed with ice-cold phosphate-buffered saline (PBS) and prepared for histology (Neuroscience Associates Inc., Knoxville, TN) [40].

### Histology

For histology, 64 rat brain hemispheres (n = 5-6/group) were drop-fixed in 4% paraformaldehyde for 24 hours, transferred to fresh PBS with sodium azide, and shipped to Neuroscience Associates (Knoxville, TN) where they were embedded into two gelatin blocks (MultiBrain_®_ Technology, NeuroScience Associates, Knoxville, TN) to be processed for histological and immunohistochemical staining. 40 μm thick sections were taken in the coronal plane, stained with de Olmos amino-cupric silver or glial fibrillary acidic protein (GFAP); primary Ab: Dako, Z0334, 1:75,000; secondary Ab: Vector, BA-1000) using the free-floating technique, visualized using 3,3’-Diaminobenzidine (DAB), and wet-mounted on 2% gelatin-subbed slides, and cover slipped. Nine photomicrographs of the S1BF and one photomicrograph of the VPM and TRN were obtained from three adjacent slides at 40× magnification on a Zeiss microscope (AXIO imager M.2). Images were captured using Neurolucida®software after the contours of anatomic landmarks taken from Paxinos and Watson [40] were outlined in each coronal section to ensure consistent locations of images. The anatomical landmarks for brain regions and whisker barrel circuit are shown in Fig. 1 and supplementary Fig. 1, respectively. GFAP: For the S1BF, 9 images per hemisphere were captured on 3 adjacent slides for a total of 27 images/rat between −2.7 mm to −3.2 mm (numbers correspond to anterior-posterior distance from bregma) using a systematic approach to represent all depths of the S1BF (See Fig. 1). In the VPM (between −2.5 mm to −3.0 mm) and TRN (between −2.5 mm to −3.0 mm) one image was obtained in each region (see Fig. 1) on 3 adjacent slides for a total of 3 images/rat. For silver stain, images were acquired from adjacent serial sections. For the S1BF, 3 images were acquired from the medial superficial, middle, and deep layers for 9 images/rat (see placement 2, 5, and 8 on Fig. 1). One image from the VPM and TRN was acquired on 3 adjacent sections for 3 images/rat/region. For the present study, the immunohistochemical staining was carried out professionally with established procedures by Neuroscience Associates Inc., and the methodology is fully validated and accurate.

**Figure 1.**
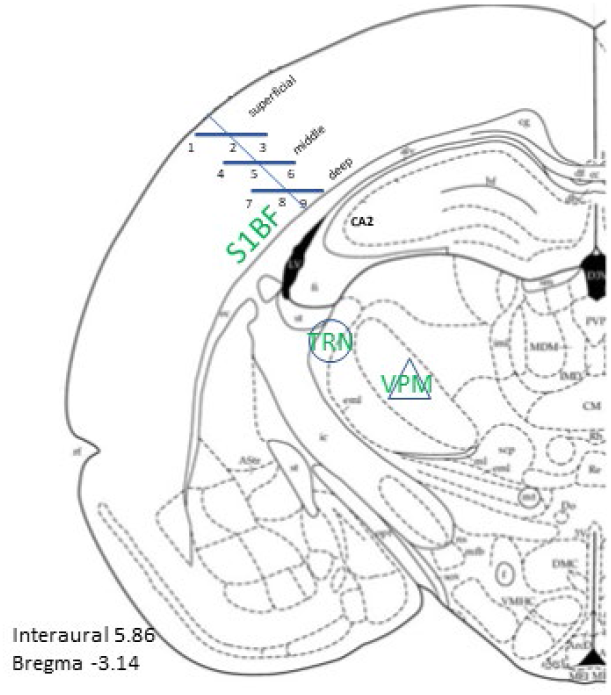
Representation of anatomical landmarks for imaging are outlined in blue. Figure obtained from Paxinos and Watson [40] showing coronal view at location bregma −3.14. S1BF region was divided into 3 divisions and images were taken medial to lateral along each line for a total of 9 images per section.

### Microscopy and analysis

#### Analysis of Amino-Cupric Silver Staining

One image was acquired in each region of the WBC for each animal (n = 5-6 per group; a total of 64 animals; a total of 192 images). Images were processed identically through ImageJ software as shown in Fig. 2. Briefly, the captured images were converted to a grayscale image with the application of eight-bit format conversion and auto-threshold function using the “Default” setting. This generated a binary image with a black signal with a white background. The “Analyze” function using the “Analyze Particles” operation in ImageJ was selected to generate a third image with size set to 0-5000 for all images (Fig. 2B and C). The final image confirms that red blood cells minimally influenced measurements (Fig. 2D). The number of pixels was calculated using the Histogram function to obtain the total number of pixels. Data were analyzed using a two-way ANOVA with Uncorrected Fisher’s LSD for comparison, as no sex effects or sex interactions were detected. Graphs represent the mean percent (%) black pixels +SEM per group.

**Figure 2.**
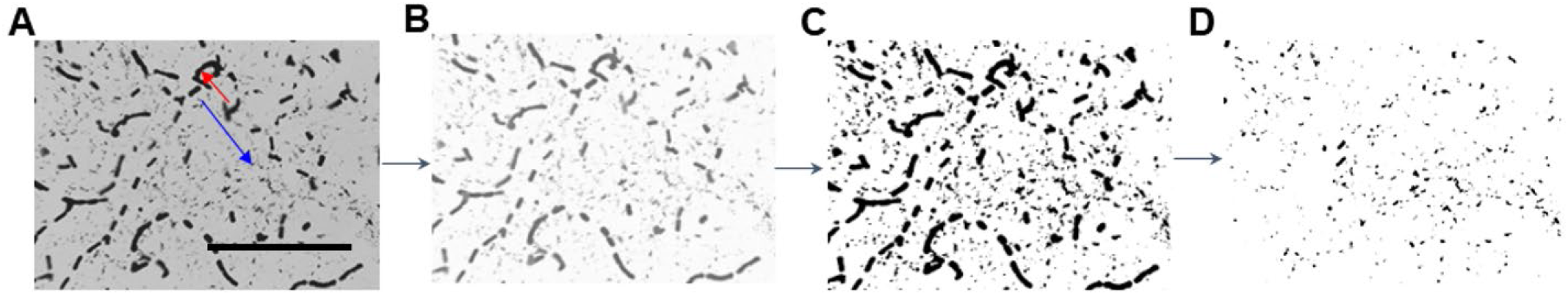
Image processing pipeline using ImageJ taken with a 40× objective. **(A)** Representative raw image of amino-cupric silver stain. Red arrow shows red blood cell staining and blue arrow is isolated neuropathology. Scale bar = 300 mm. **(B)** The raw image was converted to 8-bit file and background subtracted to give a corrected image with a black signal. **(C)** The third image represents the outlined pixels. **(D)** The number of pixels generated was calculated by using the Histogram function.

#### Analysis of Astrocyte Activation

A densitometric quantitative analysis was performed on GFAP tissue staining at 40× magnification using ImageJ software (1.48v, National Institutes of Health, Bethesda, MD, USA) employing previously published methods by an investigator with no knowledge of experimental group [32, 34, 41]. Images were converted to binary, the background was subtracted, and each image was digitally thresholded to separate positive-stained pixels from unstained pixels, then segmented into black and white pixels, indicative of positive and negative staining, respectively. The percentage of GFAP (black) staining was calculated using the following formula: [(Total area measured black/total area measured) × 100 = the percentage of area stained with GFAP]. The percentage of area stained was averaged to a single value representative of each rat per region for statistical analysis (see Fig. 3B). For the S1BF, minimal differences were identified between the 3 layers, so all 27 images were averaged to a single value per animal. Stained cells were manually counted for each image using ImageJ’s multipoint tool. Only cells with visible soma were counted. The number of cells in each image was averaged to create a single value for each animal for statistical analysis (see Fig. 3A).

**Figure 3.**
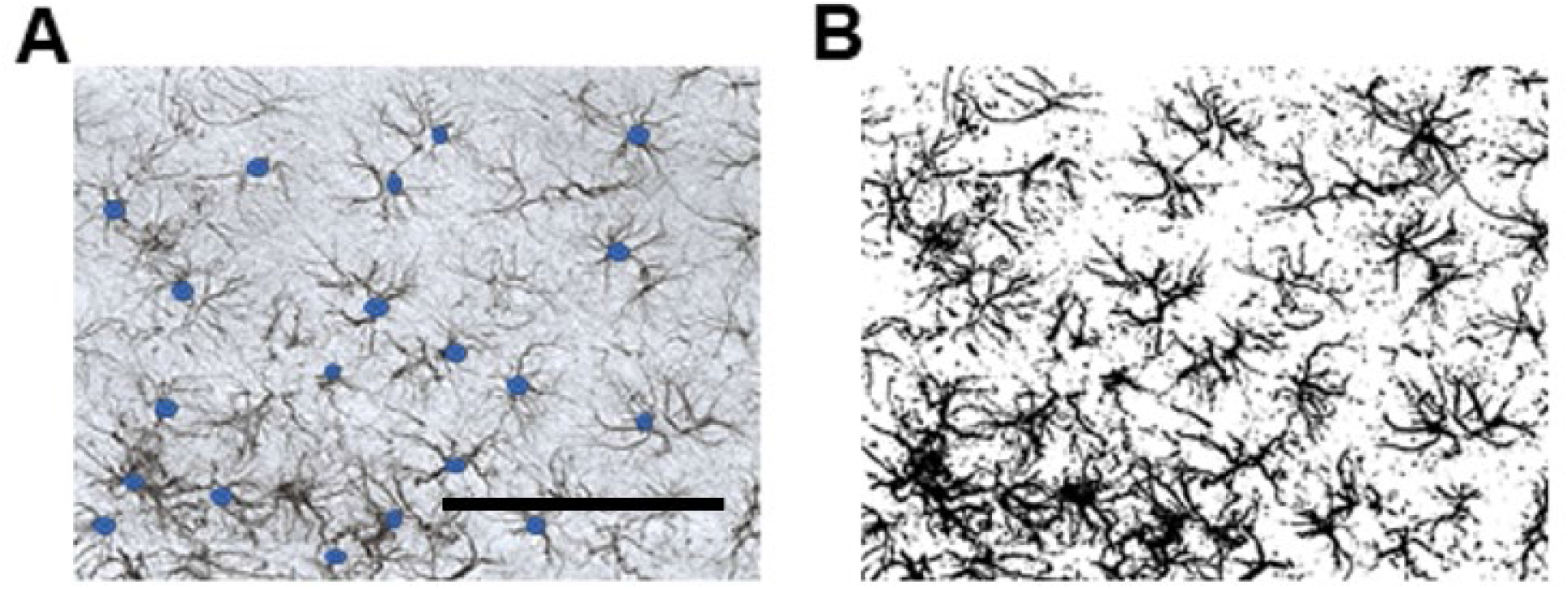
Image processing pipeline using ImageJ taken with a 40× objective. **(A)** Representative raw image immunostained for GFAP where blue dots indicate counted GFAP-positive soma with typical astrocyte morphology. **(B)** Binary image conversion in ImageJ to a black signal in a white background was used for pixel density calculations.

### Statistics

For statistical analysis, a three-way ANOVA was carried out using injury (FPI, sham), DPI (7, 56, 168), and sex (male, female) as factors. All data were tested for normality, and if data failed normality testing (Kolmogorov-Smirnov), biologically relevant data were logarithmically transformed and analyzed. Actual data (not transformed) are shown in all graphs. If no sex effects nor evidence of sex interactions were detected, a two-way ANOVA was performed on consolidated sex data. Fisher’s LSD-corrected post hoc tests were performed as appropriate to clarify all the main effects (*P* < 0.05) and interactions (*P* < 0.10). Images of astrocytes and neuropathology are from adjacent sections where data can be directly correlated. Data are presented as the mean + SEM. All statistical data were analyzed using GraphPad Prism (version 8.4.0). All data are biologically relevant in the current study, although in one region, one animal was omitted as an outlier (5% ROUT test). To fulfill ANOVA assumptions, outliers were identified and removed following logarithmic transformation if the normality test was still not passed.

## RESULTS

### VPM: Evidence of neuropathology as a function of TBI and age-with-TBI

Positive silver stain was quantified on adjacent sections as an indicator of neuropathology in male and female rats at 7, 56, and 168 DPI (Fig. 4A). No sex differences or interactions were detected (Fig. 4B), so pixel density of silver staining in male and females was consolidated for analysis as a function of FPI and DPI (Fig. 4C). Pixel density significantly increased as a function of FPI [F_(1,58)_ = 154, *P* < 0.0001] and DPI [F_(2,58)_ = 7.64, *P* < 0.01]. An interaction was observed between FPI and DPI [F_(2,58)_ = 13.6, *P* < 0.0001] (Fig. 4C), where pixel density was greater at 7 DPI (*P* < 0.0001), 56 DPI (*P* < 0.0001), and 168 DPI (*P* < 0.01) in comparison to age-matched shams. Pixel density significantly decreased between 168 DPI and 7 DPI (*P* < 0.05). In shams, pixel density increased at 56 days (*P* < 0.01) and 168 days (*P* < 0.05) compared to 7-day shams.

**Figure 4.**
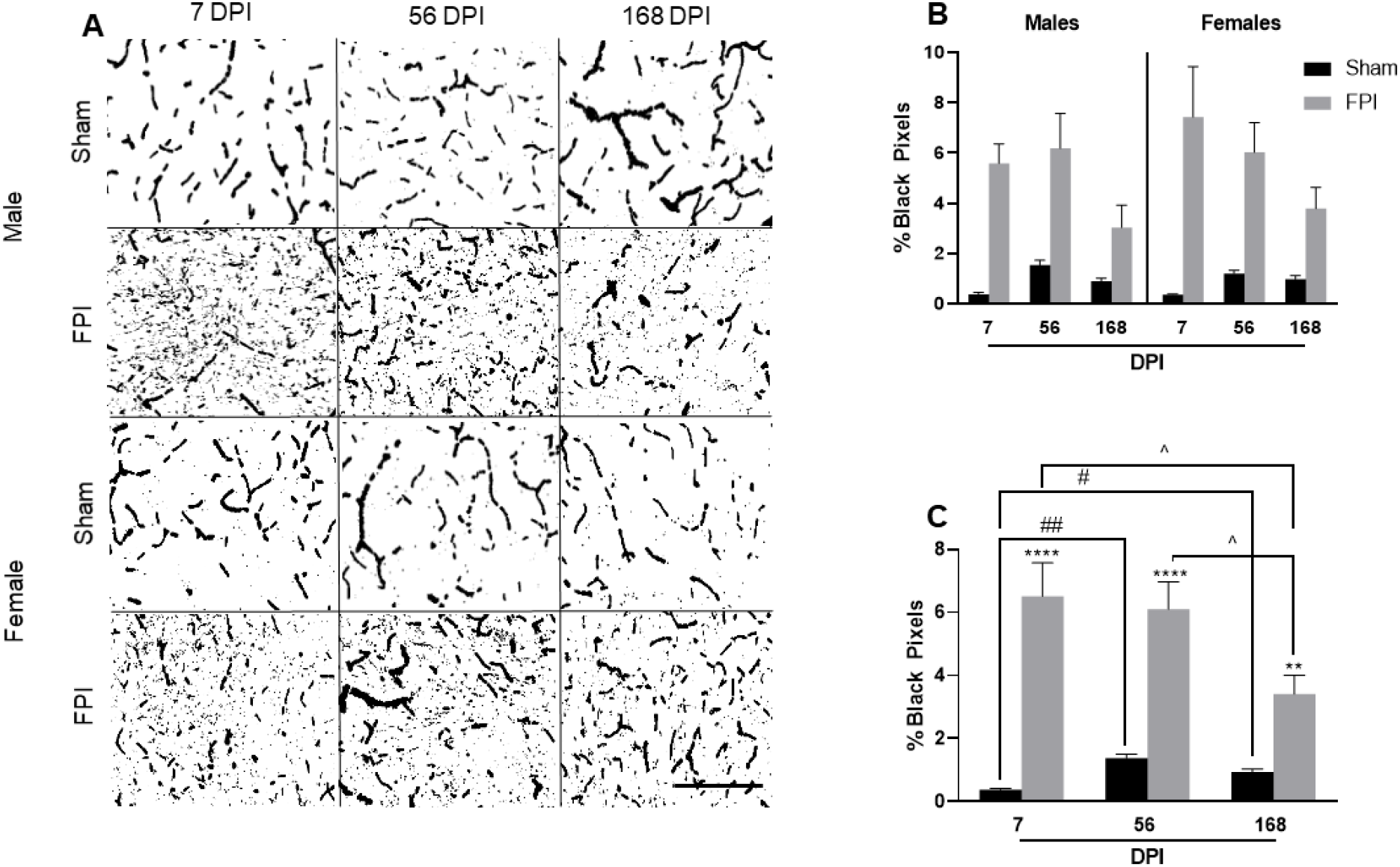
Neuropathology in the VPM as a function of aging with diffuse TBI and aging of shams. **(A)** Representative binary converted images of the VPM from sham or FPI rats of each sex stained with amino-cupric-silver technique. Scale bar = 300 μm. **(B)** Percent black pixels were quantified in VPM of each sex at 7, 56, and 168 DPI of sham and FPI rats. **(C)** Pixel density of silver staining increased as a function of FPI, DPI, and FPI × DPI interaction. Sex consolidated assessment showing diffuse TBI induced neuronal pathology at 7, 56, and 168 DPI in comparison to age-matched shams. An increase in neuronal pathology was also detected in shams at days 56 and 168 in comparison to day 7 shams. Data are expressed as the percent total black pixels (mean +SEM; *n* = 5-6 per group). \*\**P* < 0.01, \*\*\*\**P* < 0.0001, compared to mixed-sex age-matched shams; #*P* < 0.05, ##*P* <0.01, compared to mixed-sex shams at different time points; ^*P* < 0.05, compared to mixed-sex FPI at different time points.

### VPM: Chronic astrocyte activation as a function of time post-injury

As illustrated in representative photomicrographs (Fig. 5A), FPI chronically increased GFAP immunoreactivity in the VPM. There was no evidence of sex or sex interactions; therefore, the data were consolidated for a two-way ANOVA with FPI and DPI as factors (Fig. 5B and 5C). GFAP increased as a function of FPI [F_(1,58)_ = 48.53, *P* < 0.0001], DPI [F_(2,58)_ = 10.67, *P* < 0.001], and FPI × DPI interaction [F_(2,58)_ = 5.30, *P* < 0.01]. FPI increased the VPM pixel density of GFAP at 7 and 56 DPI compared to age-matched sham rats. An FPI × DPI interaction was observed, where pixel density for FPI rats increased at 7 DPI and remained at the same elevation for the time course duration. In sham controls, pixel density increased over time, reaching significance at 168 DPI (*P* < 0.001), where no difference in pixel density could be detected between sham and FPI rats at 168 DPI (Fig. 5C).

**Figure 5.**
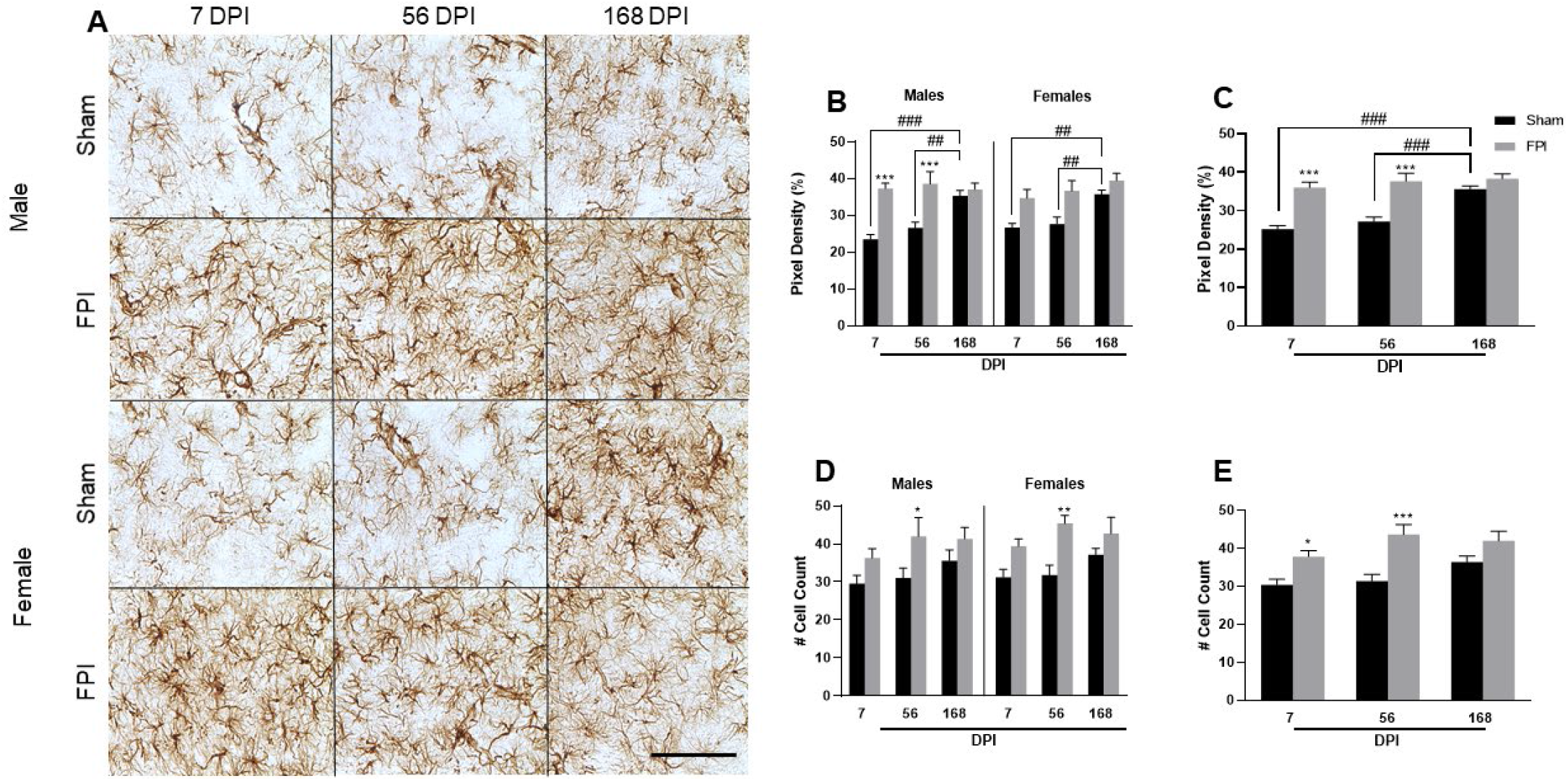
Diffuse TBI increased astrocyte activation and cell counts in the VPM at 7 and 56 DPI. **(A)** Representative photomicrographs of GFAP immunoreactivity (astrocytes) in the VPM of sham and FPI rats of each sex. Scale bar = 300 μm. **(B)** Percent black pixels quantified in VPM of sham and FPI rats of each sex at 7, 56, and 168 DPI. GFAP intensity increased as a function of FPI, DPI, and FPI × DPI interaction. Age-matched shams showed increased pixel density at day 168. **(C)** Sex consolidated percent black pixels in VPM showing increased GFAP staining at 7 DPI and 56 DPI in comparison to age-matched shams. In shams, GFAP pixel density significantly increased over time, with 168-day shams being greater than 7-day shams. At 168 DPI, no differences can be detected between sham and FPI rats. **(D)** Number of cell count in VPM of sham and FPI rats of each sex at 7, 56, and 168 DPI showed no indications of either sex differences or interactions. **(E)** Sex consolidated cell count assessment showed increased number of stained astrocytes with FPI, with significance achieved at 7 DPI and 56 DPI compared to age-matched shams. Data are expressed as mean +SEM; *n* = 5–6 per group. \**P* < 0.05, \*\**P* < 0.01, \*\*\**P* < 0.001, compared to either sex- and age-matched or mixed-sex and age-matched shams; ##*P* < 0.01, ###*P* < 0.001, compared to either sex-matched or mixed-sex shams at different time points.

To assess astrocyte proliferation, astrocyte cell somas (GFAP-positive cells) were counted in VPM. Similar to GFAP pixel density, there were no indications of sex differences or interactions (Fig. 5D), and the sexes were consolidated for statistics (Fig. 5E). GFAP-positive cells were significantly increased among FPI rats [F_(1,58)_ = 25.01, *P* < 0.0001], with cell numbers increased between 7 and 56 DPI in FPI rats (*P* < 0.05) compared to age-matched shams with the greatest magnitude difference observed at 56 DPI (Fig. 5E). Cell numbers remained elevated at 168 DPI in FPI rats, where an injury effect was lost due to the number of cells in sham rats approaching significance (*P* = 0.051) when 168-day shams were compared to 7-day shams, following a similar trajectory as GFAP pixel density. Together, these data support the observations in representative images (Fig. 5A), where GFAP increases in FPI rats that remained elevated over time and GFAP in shams increased over time. Despite an increase in cell number, there was no evidence of glial GFAP positive stained cells forming an obvious barrier indicative of scarring.

### S1BF: Evidence of neuropathology as a function of TBI and age with TBI

Positive silver stain was quantified on adjacent sections as an indicator of neuropathology (Fig. 6A). No sex differences or sex interactions were detected, so pixel density of silver staining in males and females was consolidated for analysis as a function of FPI and DPI. As shown in Fig. 6B, pixel density significantly increased as a function of FPI [F_(1,58)_ = 108, *P* < 0.0001], DPI [F_(2,58)_ = 14.9, *P* < 0.0001], and as an interaction between FPI and DPI [F_(2,58)_ = 9.39, *P* < 0.001]. Silver stain pixel density was significantly elevated at 7 DPI and 56 DPI in FPI rats (*P* < 0.0001, respectively), significantly declining at 168 DPI (*P* < 0.0001) compared to 56 DPI. The robust silver stain was measured at 56 DPI (~400% increase compared to age-matched shams). Further, at 168 DPI, there was no difference in pixel density between FPI and sham. In sham rats, there was a significant increase in pixel density between 7 and 56 DPI (*P* < 0.01).

**Figure 6.**
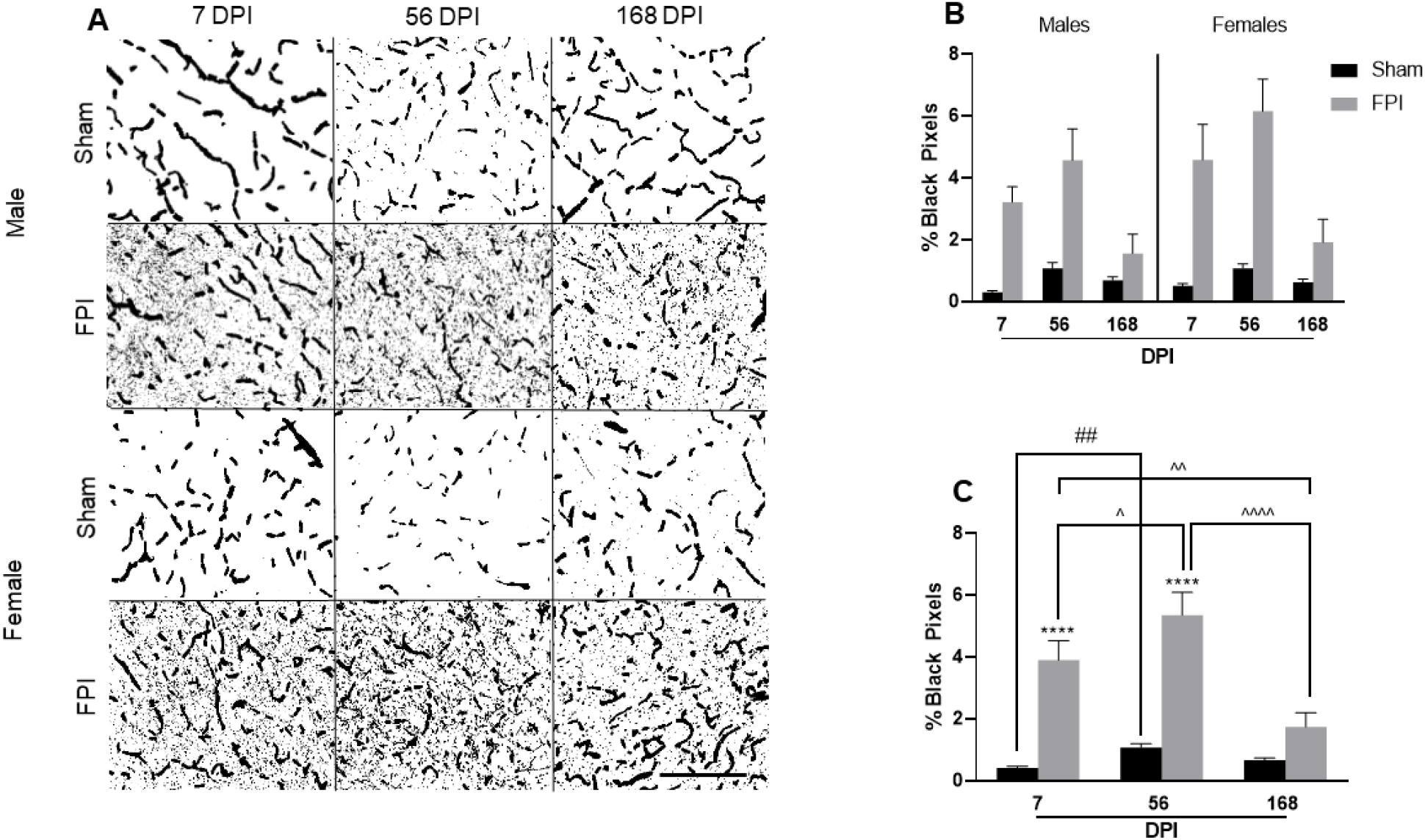
Neuropathology in the S1BF as a function of diffuse TBI and age with TBI. **(A)** Representative binary converted images of the S1BF from sham or FPI rats of each sex stained with amino-cupric-silver technique. Scale bar = 300 μm. **(B)** Percent black pixels quantified in S1BF of each sex at 7, 56, and 168 DPI of sham and FPI rats indicated no sex differences or sex interactions. **(C)** Sex consolidated assessment showed diffuse TBI-induced neuronal pathology. In particular, pixel density increased as a function FPI, DPI, and FPI × DPI interaction. Age-matched sham rats at 56 days showed increased percent of black pixels. Data are expressed as the percent total pixels (mean +SEM; *n* = 5–6 per group). *\*\*\*\*P* < 0.0001, compared to mixed-sex age-matched shams; ^##^*P* < 0.01, compared to mixed-sex sham at 7 DPI; ^*P* < 0.05, ^^*P* < 0.01, ^^^^*P* < 0.0001, compared to mixed-sex FPI at different time points.

### S1BF: Evidence of astrocyte activation as a function of time post-injury and sex

In the S1BF, GFAP immunoreactivity changed as a function of FPI [F_(1,51)_ = 99.96, *P* < 0.0001], DPI [F_(2,51)_ = 13.12, *P* < 0.0001], DPI × FPI [F_(2,51)_ = 38.40, *P* < 0.0001], and Sex × FPI [F_(1,51)_ = 3.60, *P* = 0.064]. Due to the sex interaction, data were not consolidated for the primary analysis. As demonstrated in representative photomicrographs (Fig. 7A) and pixel density quantification (Fig. 7B), FPI increased GFAP immunostaining at 7 DPI, which subsequently decreased at 56 DPI and 168 DPI compared to 7 DPI in both sexes (*P* < 0.0001). GFAP intensity was slightly higher in FPI males compared to FPI females at all 3 time points (Sex × FPI), such that FPI increased pixel density at 7 DPI (100%), 56 DPI (28%), 168 DPI (11%) in males (*vs*. age and sex-matched shams), compared to 7 DPI (83%), 56 DPI (18%), and 168 DPI (−5%) for females (*vs*. age and sex-matched shams). At 168 DPI, no differences between FPI and sham (both sexes) were detected. The FPI × DPI interaction [F_(2,58)_ = 25.23, *P* < 0.0001] can be more clearly visualized when sexes are consolidated (Fig. 7C). As pixel density decreased 35% in FPI rats between 7 and 168 DPI (*P* < 0.0001), it significantly increased by 22% in shams between 7 and 168 DPI (*P* < 0.01). One 56-day injured male was removed as an outlier (see statistics section).

**Figure 7.**
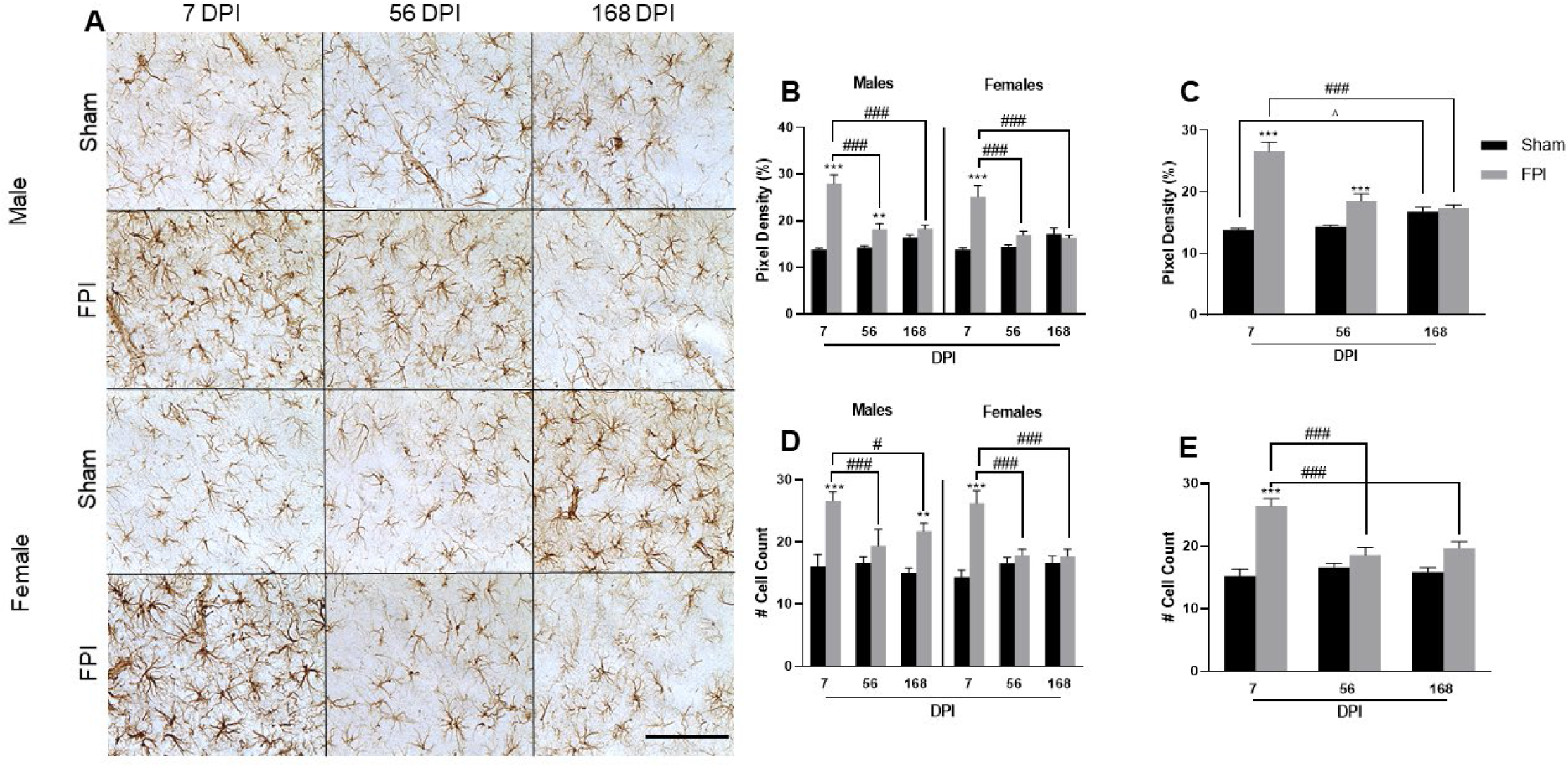
Diffuse TBI increased astrocyte activation and cell counts in the S1BF across both sexes acutely at 7 DPI. **(A)** Representative photomicrographs of GFAP immunoreactivity (astrocytes) in the S1BF of sham and FPI rats of each sex. Scale bar = 300 μm. **(B)** % black pixels quantified in S1BF of sham and FPI rats of each sex at 7, 56, and 168 DPI. GFAP immunoreactivity significantly changes as a function of FPI, DPI, DPI × FPI, and Sex × FPI. The FPI × Sex interaction was driven by overall higher GFAP immunoreactivity in males at all time points compared to females. **(C)** Sex consolidated % black pixels showed a significant FPI × DPI interaction, where pixel density decreased between 7 and 168 DPI in FPI rats and increased between 7 and 168 days in sham rats. **(D)** The astrocyte number increased with FPI, DPI, and FPI × DPI interaction. **(E)** Sex consolidated data indicated that FPI rats showed a decline in the astrocyte number over time, while the numbers in sham rats remained the same over time. Data are expressed as mean +SEM; *n* = 5–6 per group. \*\**P* < 0.01, \*\*\**P* < 0.001, compared to sex- and age-matched or mixed-sex and age-matched shams; #*P* < 0.05, ###*P* < 0.001, compared to same-sex or mixed-sex FPI at different time points; ^*P* < 0.05, compared to mixed-sex shams at 7 DPI.

GFAP immune-positive cells in S1BF also varied as a function of FPI [F_(1,51)_ = 47.45, *P* < 0.0001], DPI [F_(2,51)_ = 6.25, *P* < 0.01], and FPI × DPI interaction [F_(2,51)_ = 11.42, *P* < 0.0001]. There were no sex differences or sex interactions (Fig. 7D), and the sexes were consolidated for statistical analysis (Fig. 7E). FPI caused a 74% increase in cell numbers at 7 DPI (compared to age-matched shams) that declined by 25% as a function of DPI, where no differences were detected at 56 DPI, or 168 DPI compared to age-matched shams. A significant FPI × DPI interaction was apparent with FPI rats showing decline in cell count over time while it remained the same over time in sham rats. One 56-day injured male was removed as an outlier (see statistics section).

### TRN: Evidence of neuropathology after TBI

The TRN houses inhibitory neurons responsible for regulating sensory information in the thalamocortical circuit (VPM activity). Positive silver stain was quantified on adjacent sections as an indicator of neuropathology (Fig. 8A). No sex differences or sex interactions were detected (Fig. 8B), so pixel density of silver staining in males and females was consolidated for analysis as a function of FPI and DPI. As shown in Fig. 8C, Pixel density significantly changed as a function of FPI [F_(1,58)_ = 133.6, *P* < 0.0001], DPI [F_(2,58)_ = 6.19, *P* < 0.01], and FPI × DPI [F_(2,58)_ = 9.33, *P* < 0.001]. FPI resulted in increased silver stain pixel density a 7 DPI (*P* < 0.0001), 56 DPI (*P* < 0.0001), and 168 DPI (*P* < 0.01) in comparison to in shams, where evidence of neuropathology was present and consistent through the 168 DPI time course. In sham rats, pixel density increased as a function of time, reaching significance at 56 (*P* < 0.01) and 168 DPI (*P* < 0.05) compared to shams at 7 DPI.

**Figure 8.**
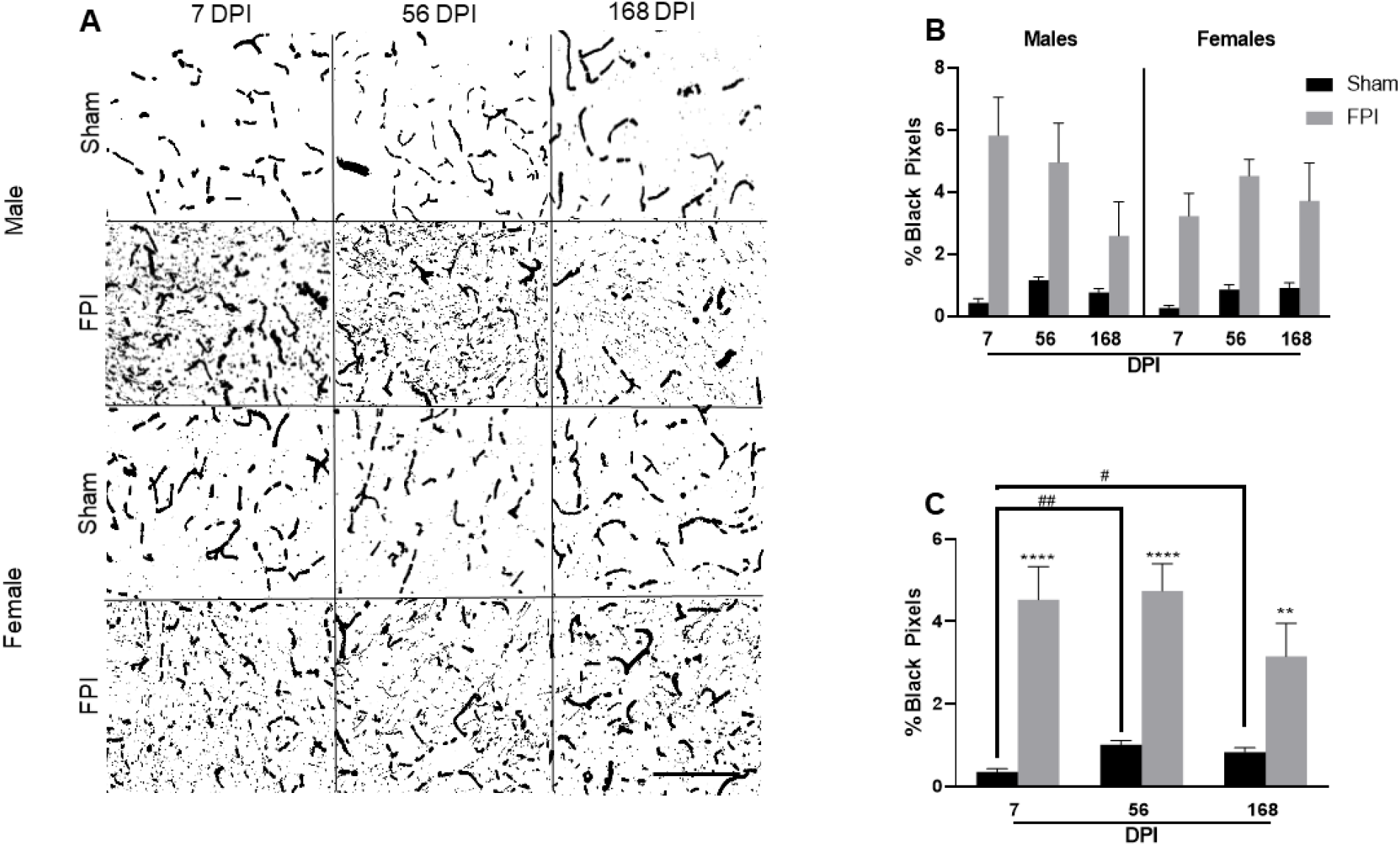
Neuropathology in the TRN as a function of diffuse TBI alone. **(A)** Representative binary converted images of the TRN from sham or FPI rats of each sex stained with amino-cupric-silver. Scale bar = 300 μm. **(B)** Percent black pixels quantified in TRN of each sex at 7, 56, and 168 DPI showed no sex differences or sex interactions in rats subjected to either sham or FPI. **(C)** Sex consolidated assessment showed changes in silver staining as a function of FPI, DPI, and FPI × DPI interaction. Increased neuropathology was evident compared to aged-matched shams at 7, 56, and 168 DPI. Data are expressed as the percent total black pixels (mean +SEM; *n* = 5-6 per group). *\*\*P* < 0.01, *\*\*\*\*P* < 0.0001, compared to mixed-sex and age-matched shams; #*P* < 0.05, ##*P* < 0.01, compared to mixed-sex shams at different time points.

### TRN: Evidence of a similar pattern of astrocyte activation as the VPM

FPI increased GFAP immunoreactivity in the TRN, demonstrated in representative photomicrographs (Fig. 9A). Since there were no indications of sex differences or interactions (Fig. 9B), the sexes were consolidated for analysis (Fig. 9C). GFAP pixel density increased as a function of FPI [F_(1,58)_ = 35.55,*P* < 0.0001], DPI [F_(2,58)_ = 23.59, *P* < 0.0001], and FPI × DPI [F_(2,58)_ = 3.14, *P* = 0.051]. Similar to the results in the VPM, GFAP pixel density increased as a function of FPI, with increased density at 7 DPI (*P* < 0.001) and 56 DPI (*P* < 0.001) in comparison to age-matched shams. As shown in Fig. 9D, in sham animals, GFAP intensity increased as a function of time, reaching significance at day 168 compared to days 7 (*P* < 0.001) and 56 (*P* < 0.001). In case of cell count in TRN, the two-way ANOVA indicated changes as a function of FPI [F_(1,52)_ = 4.33, *P* < 0.05] and DPI [F_(2,52)_ = 3.32, *P* < 0.05]. Cell counts were significantly increased at 56 DPI (*P* < 0.05) in comparison to age-matched shams. In shams, cell counts increased over time, reaching significance at day 168 in comparison to day 7 (*P* < 0.05) and 56 (*P* < 0.01).

**Figure 9.**
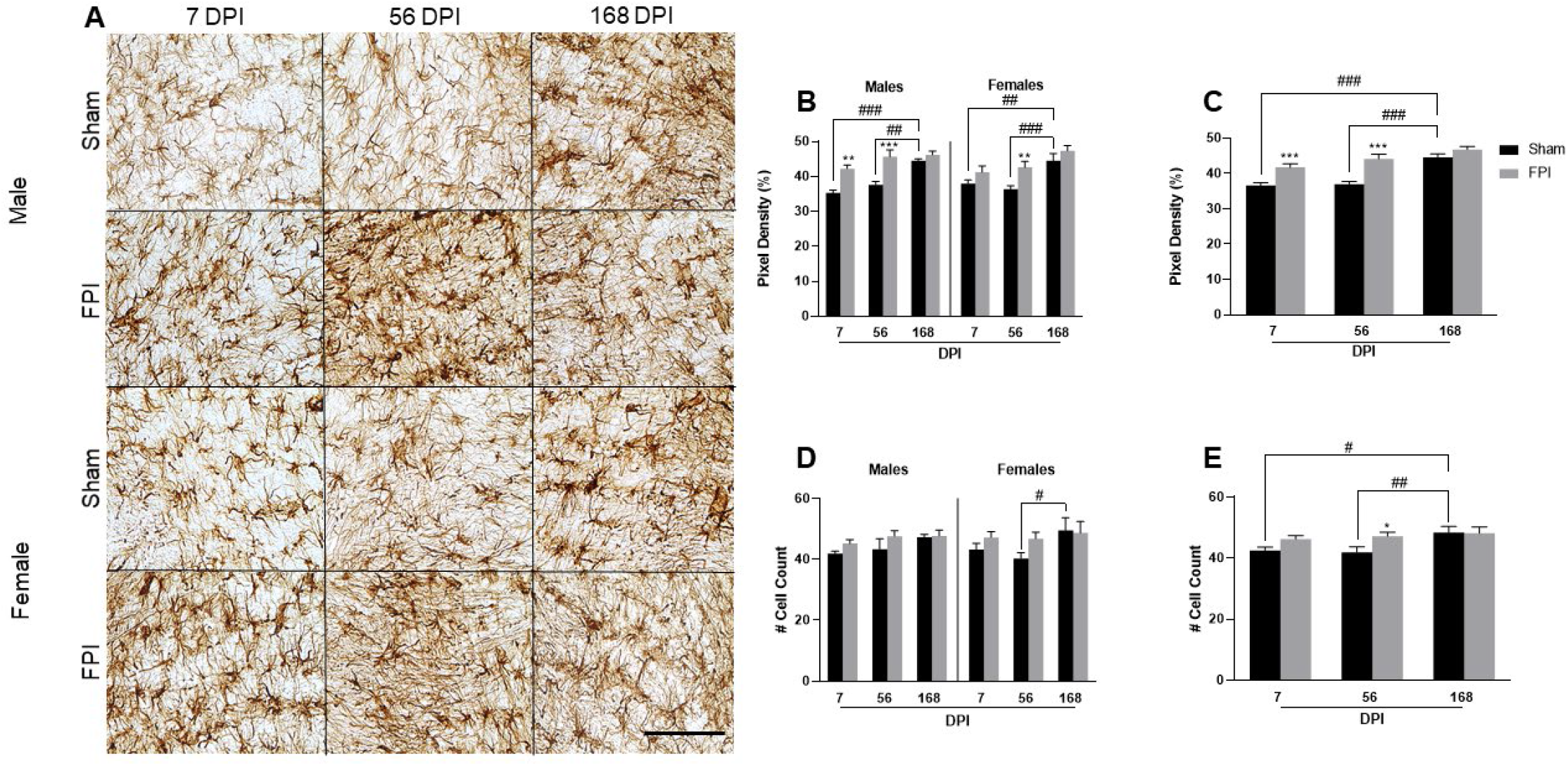
Diffuse TBI increased astrocyte activation in the TRN at 7 and 56 DPI. **(A)** Representative photomicrographs of GFAP immunoreactivity (astrocytes) in the TRN of sham and FPI rats of each sex. Scale bar = 300 μm. **(B)** Percent black pixels quantified in TRN showed increased GFAP positive density with FPI and DPI. **(C)** Sex consolidated analysis showed increased pixel density after FPI, with density being increased at 7 DPI (*P* < 0.001) and 56 DPI (*P* < 0.001) in comparison to age-matched shams. **(D)** The number of cells in TRN showed an effect of FPI on females (*P* < 0.05). **(E)** Sex consolidated analysis showed cell counts increased in shams over time, reaching significance at 168 DPI in comparison to 7 DPI (*P* < 0.05) and 56 DPI (*P* < 0.01). Data are expressed as mean +SEM; *n* = 5-6 per group.

## DISCUSSION

The WBC has high circuit integrity of relays for each whisker (owing to its somatotopographical organization) for the flow of sensory information [29, 42]. In the thalamic relay (i.e. VPM) of the WBC, there are few interneurons and inhibition comes from GABA projections via the TRN and modulation from NE projections from the locus coeruleus [29]. Previous studies have indicated that the WBC is vulnerable to axonal, glial, and vascular disruption during and in the aftermath of experimental TBI contributing to the development of late-onset and persisting sensory hypersensitivity [28, 43, 44]. Along these lines, experimental TBI in the WBC provides an *in vivo* model of diffuse axonal injury with fewer innervating circuits, neurotransmitter types, and interneurons, minimizing confounds by which we can: (1) gain knowledge about the benefits and detriments of temporal pathophysiological processes responsible for the development or persistence of morbidity, (2) understand the impact of interventions on these processes in mediating TBI-related morbidity, and (3) assess mechanisms associated with increased risks for age-related diseases [30, 45, 46].

Our results indicate neuropathology and astrogliosis are increased as a function of injury and remain elevated to 168 DPI in the thalamic nuclei. In the S1BF, both pathologies are increased at 7 DPI and tend to decline over time. However, they are still significantly elevated at 56 DPI and achieve congruency with shams by 168 DPI. Shams in all regions showed evidence of age-related neuropathology and increasing astrocyte immunoreactivity at 56 and/or 168 days post-sham procedures in comparison to 7-day shams, indicating that the similarity between sham and brain-injured rats at 168 DPI is likely driven by a convergence of age-related pathology with TBI-related pathology. These data indicate dynamic time-, region-, and sex-dependent pathological changes that are likely pertinent to the development and persistence of whisker hypersensitivity. These experiments support the utility of using the WBC, specifically the thalamocortical relay, to study the divergence of cellular and molecular phenotypes responsible for TBI-induced morbidity and the potential to evaluate the convergence of neuropathology and astrogliosis as a function of age and aging with TBI with translational relevance to higher-order networks. Further, we can translate these data to assist in interpretation of pathophysiological data from more complex circuitry, like the amygdala or hypothalamus, which can be confounded due to an exceptionally high degree of innervations from other brain regions and white matter tracts injury after TBI [14, 31, 32, 34, 42].

### Chronic Neuropathology Following a Single TBI

Amino-cupric silver stain for damaged and degenerating neurons [47] indicated acute neuropathology with varying degrees of sustained neurodegeneration across all time points and regions after TBI. The sustained neuropathology observed in VPM and TRN up to 168 DPI is likely associated with axotomy and Wallerian degeneration in ascending and descending fiber tracts resulting in argyrophilic accumulation, but could also be perpetuated by secondary injury cascades leaving the thalamus more vulnerable to late-onset axonal damage or functional impairment [48, 49]. Thalamic axonal damage associated with gliosis has been demonstrated out to 28 DPI in male brain-injured rats [33], supported by post-mortem human pathology indicating thalamic (and cortical) atrophy as common features of chronic traumatic encephalopathy [50], also predominantly evaluated in males. Despite other pathological features of axonal damage being validated in humans, the presence and duration of Wallerian degeneration in chronic TBI patients has not been extensively evaluated, where our data indicate the thalamus may be more vulnerable to chronic neuropathology that can only be detected using this or a similar method. Although scant, some previous reports indicate selective neuronal loss of GABAergic cells and processes in the TRN and VPM of individuals after severe trauma and in preclinical models using lateral FPI, respectively [51, 52]. Our results indicate that TRN projections and GABA innervation may be disrupted chronically after experimental mild TBI. Increased thalamic axotomy associated with neuronal atrophy and reconnection (not cell death) within glutamatergic and GABAergic projections promotes an environment for global circuit dysfunction, likely underlying whisker hypersensitivity [33, 53]. Accumulating evidence also supports a role for chronic TBI-induced thalamic connectivity issues with sleep disruption, attention, executive control, perceptual decision-making, headache/migraine, and other complex behaviors mediated by coordinated thalamic activity, where the thalamus may play a key role in symptom development and recovery after TBI [54–57].

Region-dependent changes in neuropathology are likely multifactorial, where the time-dependent decrease in the S1BF may also be secondary to the gradual progression of argyrophilic reaction product from superficial to deep cortical layers due to axonal damage to corticothalamic and thalamocortical projections coupled with complex connectivity between cortical layers [44, 58]. Doust et al. reported increased evidence of amyloid precursor protein and SM134 pathology in the S1BF despite age at injury (PND17-6mos) and aging with injury (4-10 months) in comparison to 10-month-old naïve controls in males using midline FPI, with the S1BF and hippocampus being most prone to chronic pathology [59]. While this study focused on axonal pathology of intact axons, it does support the notion that the S1BF is vulnerable to chronic neuropathology beyond Wallerian degeneration. Overall, these data are consistent with the accumulating evidence of long-lasting neuronal pathology in the somatosensory (and other) relays after a diffuse TBI, but extend these observations to six months post-injury and in females [29, 33, 59–62]. The presence of TBI induced neuropathology out to 6 months post-injury in an experimental model of TBI indicates ongoing Wallerian degeneration which may coincide with characteristic late CTE pathology (tau and amyloid pathologies) found at 10-20 years post-injury [63]. Overall, these data support that a single diffuse TBI can lead to long-term neurodegeneration, with increased vulnerability of relay structures, like the thalamus, further supporting that TBI instigates neurodegenerative disease.

### Chronic Astrogliosis Following a Single TBI

In addition to axonal degeneration, TBI disrupts overall brain homeostasis eliciting chronic astrogliosis, where GFAP immunoreactivity and cell numbers were significantly increased at 7 DPI in all regions. Astrogliosis is characterized by functional, morphological, and molecular remodelling, in addition to the release of immunoregulatory factors which may promote or limit inflammation and post-injury recovery processes [7, 16, 23]. Previous reports indicate that TBI increased macrophage infiltration and microglial activation to facilitate clearance of degenerating neurons over time, promote the neuroinflammatory response, or alternatively, promote synaptic and neuronal reorganization [33, 64–67]. Reactive microglia and macrophages can directly activate astrocytes, as can the massive disruption in homeostasis (altered circuit function, metabolic crisis, ion balance, and excess degenerating neurons). Predominantly, astrocytes are implicated in promoting recovery and regaining homeostasis by promoting cellular homeostasis and mediating neuroplastic events [7, 44, 68]. Along these lines, the astrogliosis may provoke axonal sprouting and new synaptic connections of adaptive and maladaptive reorganization, where maladaptive is implicated in development of behavioral morbidity like late-onset sensory hypersensitivity to whisker stimulation which has been replicated across several labs [14, 27, 28, 43, 69]. It is likely that mechanisms mediating preclinical morbidity are relevant to clinical reports where TBI-induced damage to the thalamus is associated with sensory hypersensitivity and hyperarousal which leads to agitation and aggressive behavior in brain-injured humans [70, 71]. One potential mechanism for activated astrocytes to mediate post-injury plasticity is that they are a source for thrombospondins and other synaptogenic molecules, implicating hyperconnectivity of glutamatergic neurons associated with hyperexcitability and the development of epileptiform signaling after acquired brain injury [72, 73]. Astrogliosis can also influence neurotransmission, where astrocytic glutamate transporters are responsible for the uptake of synaptic glutamate to maintain optimal extracellular glutamate levels [74]. Our own work has revealed region and sex-dependent changes in evoked glutamate release and clearance kinetics in the thalamocortical relays chronically post-TBI, where astrogliosis may play a role in neuroplasticity/neurotransmission, or a combination of both. Others have implicated astrocyte control of GABAergic inhibition in the whisker barrel regions as the major inhibitory neurotransmitter for the corticothalamic circuit [75]. Thus, our findings support roles where astrocyte disruption could influence the excitatory/inhibitory balance by which post-TBI symptoms persist. GFAP immunoreactivity and cell numbers increased at 7 DPI in VPM and TRN and remained at a similar level for the duration of the 168 DPI time course. Significance in comparison to sham was only maintained until 56 DPI, where gliosis and cell counts in shams began to increase to similar levels of brain-injured rats at 168 DPI. This observation coincided with the appearance of mild neuropathology in shams. Chronic neuropathology and delayed reactive astrogliosis in the thalamus are specifically implicated in sleep disruption, persisting TBI symptoms, accelerating aging, and increasing risk for age-related neurodegenerative diseases, where the role of astrocytes are the least understood, despite their importance in homeostasis, neuroplasticity, neurotoxicity, neurotransmission, and BBB maintenance [76, 77].

This is the first documentation of a convergence between TBI-induced astrogliosis and age-related astrogliosis. TBI instigates secondary injury cascades that are implicated in delayed cell death, and recent evidence suggests that the activated astrocytes can directly induce neuronal death [15, 78]. Mild TBI leads to increased risk for long-term decline related to ongoing and delayed neurodegeneration [79]. Aging with TBI and normal aging itself could exacerbate or accelerate pathophysiological processes to disturb neural circuit function observed both clinically [8, 80] and in experimental models [81, 82]. Further, the pathological nuances between aging with TBI and aging are not yet well understood, making it challenging to identify the pathophysiology responsible for long-term complications. The senescent astrocyte may also contribute vulnerability to compromised neurons by promoting an inflammatory response after TBI [24, 83], a mechanism which can increase risk of age-related neurodegenerative diseases. Astrocyte hypertrophy in the uninjured shams may be related to the aging process where previous studies have indicated that general brain aging itself exacerbates neuroinflammatory response [84]. For instance, immune-inflammatory hypotheses for aging have emerged, including microglia-astrocyte interactions involving microglia-driven astrocyte activation in aging and inflammation stemming from A1 reactive astrocytes [22]. It might be that this ‘inflammaging’ process affects astrocyte function and promotes secondary brain damage when such an effect is severe or prolonged. Aside from this, the astrocyte research community is evolving away from the aforementioned binary division of reactive astrocytes (A1/A2) towards a classification system with multiple molecular markers to better represent unique functional reactive astrocyte phenotypes [7]. Regardless, as aging is the primary risk factor for neurodegenerative conditions, it is crucial to evaluate the impact of TBI not in isolation, but over time to understand better the evolution of brain pathology and translate inflammatory events occurring in the patient population. The time course of this convergence could serve as a meaningful approach to evaluate phenotypic astrocytic differences after prolonged gliosis versus newly onset gliosis, with novel insights into the role of astrocytes in TBI, Alzheimer’s and Parkinson’s disease pathology.

Interestingly, the post-TBI time course of VPM/TRN astrogliosis was dissimilar to S1BF astrogliosis. In the S1BF, GFAP immunoreactivity increased by 92% at 7 DPI, falling to 30% at 56 DPI, with no difference observed at 168 DPI. The decrease between 7 and 56 DPI was not associated with a change in neuropathology, which significantly decreased between 56 and 168 DPI. These data indicate that the temporal profile of neuropathology and astrogliosis greatly differ between regions, where overall pathology in the cortex demonstrates greater propensity to return to preinjury status. The region dependence could be driven by ongoing secondary cascades responsible for slowly progressive Wallerian-like degeneration in the thalamic nuclei [85], where positive correlations between silver and GFAP pixel density were predominant in the 56 and 168 DPI thalamic nuclei compared to the S1BF (see supplemental Tables S2 and S3). This observation likely supports that astrogliosis is also region and time dependent, where neuropathology is not necessarily the driver of astrogliosis.

### Sex Differences

While previous studies have indicated sex differences in chronic temporal neuroinflammatory trajectories after controlled cortical impact [86], few have evaluated these changes in association to neuropathology and neurodegeneration in both sexes after diffuse TBI. In fact, studies have reported sex differences in inflammatory response after experimental TBI, where increased acute and subacute neuronal death and astrogliosis was greater in males compared to females, however, chronic time points demonstrated similar pathology between the sexes [31]. In the S1BF, male GFAP levels returned to sham levels at 168 DPI, and cell counts remained increased in the males out to 168 DPI, while female GFAP levels and cell counts returned to baseline levels at 56 DPI. Some evidence suggests that estrogen can regulate glial activation, involving the estrogenic receptor subtypes expressed on glia [87–89]. Moreover, it was recently demonstrated that estrogen is reported to inhibit Toll-like receptor 4 (TLR4)/nuclear factor kappa B (NF-kB) pathway-regulated astrocyte activation after TBI [90], suggesting that the sex effect stemmed from estrogen-mediated differences in GFAP upregulation in the S1BF. Other reports have documented that changes in ovarian hormones can promote changes in astroglial-mediated plasticity [91]. Consistent with our previous finding, increased evoked S1BF glutamate release was present in males at 28 DPI, but not females [14, 28]. Females also had lower overall whisker nuisance scores than males at 28 DPI, which could possibly be related to decreased evoked release or alternatively decreased intensity of GFAP mediated by estrogen-glial interactions. However, further investigation is warranted to substantiate this possibility.

## LIMITATIONS AND CONCLUSION

We fully acknowledge the limitations of restricting to one stain to assess neuropathology and astrocyte reactivity, and it is not known whether parallel changes occurred with respect to other pathological markers. While GFAP is the most widely used marker of reactive astrocytes and intensity of GFAP immunoreactivity often parallels injury severity, a more complete temporal profile including multiple astrocytic targets is still necessary to determine astrocyte phenotypes [7]. Further, assessment of other neuroinflammatory cells would enhance the interpretation, by assisting in identifying functional roles of activated astrocytes [22] and mechanisms by which different phenotypic profiles can impact how aging with TBI influences long-term recovery. Further work is needed to determine mechanisms mediating differences as a function of time post-injury, aging-with-injury, region, and sex. We did not include a naïve group in the study, where differences in the sham group could be influenced by surgery, anesthesia, or other procedures. In the current study, we did not track the estrous cycle to avoid confounds associated with stress arising from daily handling. We have previously published that FPI can decrease time spent in estrus phase [14], so it is possible that dynamic variations in the estrous cycle and circulating ovarian hormones could influence astrocyte activation (and function).

These caveats notwithstanding, our results clearly indicate distinct region-dependent chronic temporal profiles of neuropathology and astrocyte reactivity in relays of the whisker barrel circuit of males and females after a diffuse TBI. The divergent pattern of pathological outcomes observed in both sexes after brain injury is likely pertinent to the increased risk for patients to develop age-related neurodegenerative diseases. The findings indicate that thalamic relays may be at higher risk for chronic neuropathology and, thereby, physiological mechanisms essential for return to homeostasis. The present findings support the relevance of assessing pathophysiological processes to identify time-, region-, and sex-dependent pathology responsible for the development and persistence of post-TBI morbidities. Further, this information supports that TBI increases the risk for neurodegenerative disease where outcomes can serve as biomarkers in future preclinical experiments to determine the efficacy of therapies targeting chronic neurodegeneration and astrogliosis that are necessary to improve clinical diagnosis and prognosis.

## FUNDING

This work was supported by National Institutes of Health (Grant reference number R01NS100793), Arizona Biomedical Research Commission through Arizona Department of Health Services (ADHS14-00003606), Phoenix Children’s Hospital Leadership Circle Grants, and Phoenix Children’s Hospital Mission Support to TCT.

## ACKNOWLEDGMENTS

We thank Jim Baun and his team at Neuroscience Associates for assistance in generating the histology archive. We acknowledge the helpful feedback from Chaitanya Sanghadia on these data. Lastly, we thank Dr. Rachel Rowe for assistance with tissue generation. We also thank Carol Haussler for assistance with critical review of the manuscript.

## AUTHOR CONTRIBUTIONS

Zackary Sabetta, Conceptualization, Data curation, Formal analysis, Validation, Investigation, Visualization, Methodology, Writing - original draft, Writing - review and editing; Gokul Krishna, Conceptualization, Formal analysis, Writing - original draft, Writing - review and editing; Tala Curry, Writing - review and editing; David Adelson, Writing – review and editing; Theresa Currier Thomas, Conceptualization, Resources, Data curation, Formal analysis, Supervision, Funding acquisition, Investigation, Visualization, Methodology, Writing - original draft, Project administration, Writing - review and editing.

## DISCLOSURE STATEMENT

The authors declare no competing interests. This work is solely the responsibility of the authors and does not necessarily represent the official views of the funding agencies. The funders had no role in study design, data collection and interpretation, or the decision to submit the work for publication.

## Ethics

Animal experimentation: All procedures were conducted in accordance with the NIH Guide for the Care and Use of Laboratory Animals and were approved by the Institutional Animal Care and Use Committee Protocol (# 13-384) at the University of Arizona College of Medicine-Phoenix.

## Data Availability Statement

The datasets generated for this study are available upon reasonable request to the corresponding author.

## Notes

### Competing Interest Statement

The authors have declared no competing interest.

