## Supplemental Data 1 for "Aging with TBI vs. Aging: 6-month temporal profiles for neuropathology and astrocyte activation converge in behaviorally relevant thalamocortical circuitry of male and female rats"

**Supplemental Material:**
Supplemental Tables: 3
Supplemental Figures: 1

| | S1BF | | |
|---|---|---|---|
| | Silver stain vs. GFAP | Silver stain vs. Cell count | Silver stain vs. Righting reflex time (s) |
| 7 Males | ns | ns | ns |
| 7 Females | ns | ns | ns |
| 56 Males | ns | ns | ns |
| 56 Females | 0.0247 | ns | ns |
| 168 Males | ns | ns | ns |
| 168 Females | ns | 0.0464 | ns |

**Table S1. Correlation analysis of S1BF neuropathology with astrocyte activation and neuropathology with righting reflex for FPI rats.** Outcome measures include silver stain, cell counts, GFAP level, and righting reflex time in seconds. There is a significant correlation between neuropathology and cell counts at 168 DPI in female rats and between neuropathology and GFAP levels at 56 DPI in female rats.

| | VPM | | |
|---|---|---|---|
| | Silver stain vs. GFAP | Silver stain vs. Cell count | Silver stain vs. Righting reflex time (s) |
| 7 Males | ns | ns | ns |
| 7 Females | ns | ns | ns |
| 56 Males | 0.0045 | ns | ns |
| 56 Females | ns | ns | ns |
| 168 Males | 0.0263 | 0.0172 | ns |
| 168 Females | 0.0327 | ns | ns |

**Table S2. Correlation analysis of VPM neuropathology with astrocyte activation and neuropathology with righting reflex for FPI rats. Outcome measures include silver stain, cell counts, GFAP level, and righting reflex time.** There is a significant correlation between neuropathology and GFAP at 56 DPI in males, 168 DPI in males, and 168 DPI in females. A correlation was only detected at 168 DPI in males for neuropathology and cell counts.

| | TRN | | |
|---|---|---|---|
| | Silver stain vs. GFAP | Silver stain vs. Cell count | Silver stain vs. Righting reflex time (s) |
| 7 Males | ns | ns | ns |
| 7 Females | 0.0213 | ns | 0.0015 |
| 56 Males | 0.0419 | ns | ns |
| 56 Females | ns | ns | ns |
| 168 Males | ns | 0.0172 | ns |
| 168 Females | ns | ns | ns |

**Table S3. Correlation analysis of TRN neuropathology with astrocyte activation and neuropathology with righting reflex for FPI rats. Outcome measures include silver stain, cell counts, GFAP level, and righting reflex time.** There is a significant correlation between neuropathology and GFAP 7 DPI in females and 56 DPI in males, neuropathology and cell counts at 168 DPI in males, and neuropathology and righting reflex time at 7 DPI in females.

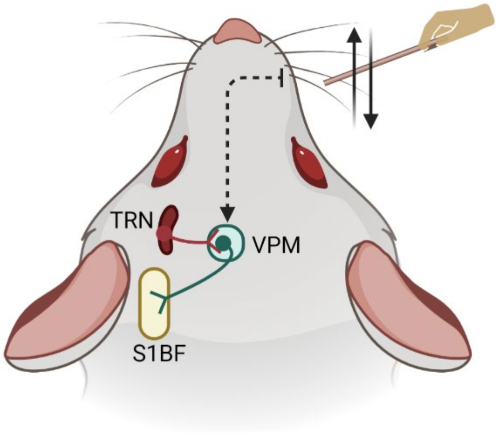


**Supplement Figure 1. Representative layout of somatosensation through the rat whisker barrel circuit.** Forward and backward movement of whiskers evokes sequence of events in trigeminal neurons following which glutamatergic projections relay through VPM (green) for higher level processing in the S1BF (yellow). The VPM is inhibited by GABAergic projections from the TRN (red). Created with BioRender.com
